# Microzooplankton distribution in the Amundsen Sea Polynya (Antarctica) during an extensive *Phaeocystis antarctica* bloom

**DOI:** 10.1101/271635

**Authors:** Rasmus Swalethorp, Julie Dinasquet, Ramiro Logares, Stefan Bertilsson, Sanne Kjellerup, Anders K. Krabberød, Per-Olav Moksnes, Torkel G. Nielsen, Lasse Riemann

**Affiliations:** Scripps Institution of Oceanography, University of California San Diego, USA; National Institute of Aquatic Resources (DTU Aqua), Technical University of Denmark, Denmark; Department of Marine Sciences, University of Gothenburg, Sweden; Marine Biological Section, Department of Biology, University of Copenhagen, Denmark; Department of Natural Sciences, Linnaeus University, Sweden; Institute of Marine Sciences (ICM), CSIC, Spain; Department of Ecology and Genetics: Limnology and Science for Life Laboratory, Uppsala University, Sweden; Department of Biosciences, Section for Genetics and Evolutionary Biology (Evogene), University of Oslo, Norway

**Keywords:** ciliate, dinoflagellate, growth rates, Southern Ocean, Antarctica, Amundsen Sea polynya, *Gymnodinium* spp.

## Abstract

In Antarctica, summer is a time of extreme environmental shifts resulting in large coastal phytoplankton blooms fueling the food web. Despite the importance of the microbial loop in remineralizing biomass from primary production, studies of how microzooplankton communities respond to such blooms in the Southern Ocean are rather scarce. Microzooplankton (ciliates and dinoflagellates) communities were investigated combining microscopy and 18S rRNA sequencing analyses in the Amundsen Sea Polynya during an extensive summer bloom of *Phaeocystis antarctica*. The succession of microzooplankton was further assessed during a 15-day induced bloom microcosm experiment. Dinoflagellates accounted for up to 58% the microzooplankton biomass *in situ* with *Gymnodinium* spp., *Protoperidium* spp. and *Gyrodinium* spp. constituting 87% of the dinoflagellate biomass. *Strombilidium* spp., *Strombidium* spp. and tintinids represented 90% of the ciliates biomass. *Gymnodinium*, *Gyrodinium* and tintinnids are known grazers of *Phaeocystis,* suggesting that this prymnesiophyte selected for the key microzooplankton taxa. Availability of other potential prey, such as diatoms, heterotrophic nanoflagellates and bacteria, also correlated to changes in microzooplankton community structure. Overall, both heterotrophy and mixotrophy appeared to be key trophic strategies of the dominant microzooplankton observed, suggesting that they influence carbon flow in the microbial food web through top-down control on the phytoplankton community.

## 1. Introduction

The Southern Ocean (SO) plays a central role in global biogeochemical cycles due to strong summer pulses of primary production (Sarmiento et al., 2004). Coastal polynyas contribute strongly to the efficiency of the biological pump through massive export of organic material (DiTullio et al., 2000) and formation of deep water masses (Williams et al., 2007). The summers, with elevated irradiance, reduced ice cover and subsequent input of nutrients, sparks a short but massive burst of phytoplankton in these areas (Smith Jr & Gordon, 1997; Arrigo et al., 2012). To understand and predict the extent of CO_2_ sequestration in the SO (Sabine et al., 2004; Arrigo et al., 2008) it is therefore important to determine the fate of the extensive phytoplankton blooms occurring in the Antarctic polynyas.

Microzooplankton have been estimated to graze over half of the daily global planktonic primary production (Calbet & Landry, 2004; Schmoker et al., 2013) and may thus exert significant top-down control on phytoplankton blooms in the SO (Bjørnsen & Kuparinen, 1991; Kuparinen & Bjornsen, 1992). Despite the key ecological functions of microzooplankton in the carbon cycle as either primary producers or as a trophic link between the microbial loop and higher trophic levels (reviewed in Sherr & Sherr, 2009), they have received little attention in the productive polynyas of the SO. The Amundsen Sea Polynya (ASP) is one of the most productive polynya of the SO, characterized by dramatic perennial blooms (Arrigo & van Dijken, 2003). In summer 2010-2011 the Amundsen Sea Polynya Research Expedition (ASPIRE) aimed to determine the fate of the high algal productivity. At the time of sampling, the ASP was undergoing an extraordinary bloom event dominated by the prymnesiophyte algae *Phaeocystis antarctica* (Alderkamp et al., 2015; Yager et al., 2016).

Microzooplankton grazing pressure on *Phaeocystis* depends on whether the prymnesiophyte occurs in its single cell or colonial form (Caron et al., 2000; Grattepanche et al., 2010; Grattepanche et al., 2011). Thus, the succession of the bloom affects the microzooplankton community, its grazing pressure (Verity, 2000), and impact on the biological pump. In the ASP, carbon export was high (up to 62% of net primary production - NPP) and grazing rates were low (32.5% of the NPP in the upper 100 m of the water column (Yager et al., 2016)) compared to other regions of the SO (Froneman & Perissinoto, 1996; Landry et al., 2002; Pearce et al., 2008; Schmoker et al., 2013). Nevertheless, microzooplankton grazing pressure on the NPP was far more important than that of mesozooplankton (Wilson et al., 2015; Yager et al., 2016) and became increasingly important later during the bloom (Lee et al., 2013; Yang et al., 2016). This underlines the importance of understanding the coupling between the microzooplankton community and these extraordinary blooms to better predict the fate of the carbon. This is pertinent as the Amundsen Sea ice-sheet and sea ice are rapidly melting due to environmental warming (Pritchard et al., 2009; Stammerjohn et al., 2015), which may modify the magnitude as well as the temporal and spatial dynamics of *P. antarctica* and diatoms blooms (Alderkamp et al., 2012).

In the present study, we examined the composition of the microzooplankton community in the ASP during an intense *P. antarctica* bloom. To improve taxonomic resolution we combined microscopy and 18S rRNA sequencing to characterize the community. Additionally, the microzooplankton succession during an intense bloom event was investigated through a 15-day induced *P. antarctica* bloom microcosm experiment. The aim was to improve our understanding of the environmental drivers shaping the community of these principal phytoplankton grazers within Antarctic polynyas during the productive Austral summer.

## 2. Materials and Methods

### 2.1. In situ conditions

The sampling was conducted during the ASPIRE cruise in November 2010 to January 2011 onboard RVIB N.B. Palmer (cruise NBP1005; (Yager et al., 2012; Yager et al., 2016). Fourteen stations were sampled for microzooplankton analyses via microscopy and 18S rRNA amplicon sequencing (Fig. 1). Water was collected with 12 l Niskin bottles mounted on a CTD rosette (Sea-Bird 911+). Detailed information on methods and *in situ* environmental parameters measured (phytoplankton pigments, nutrients, bacteria and heterotrophic nanoflagellates – HNF abundance) can be found in Yager et al. (2016).

**Figure 1:**
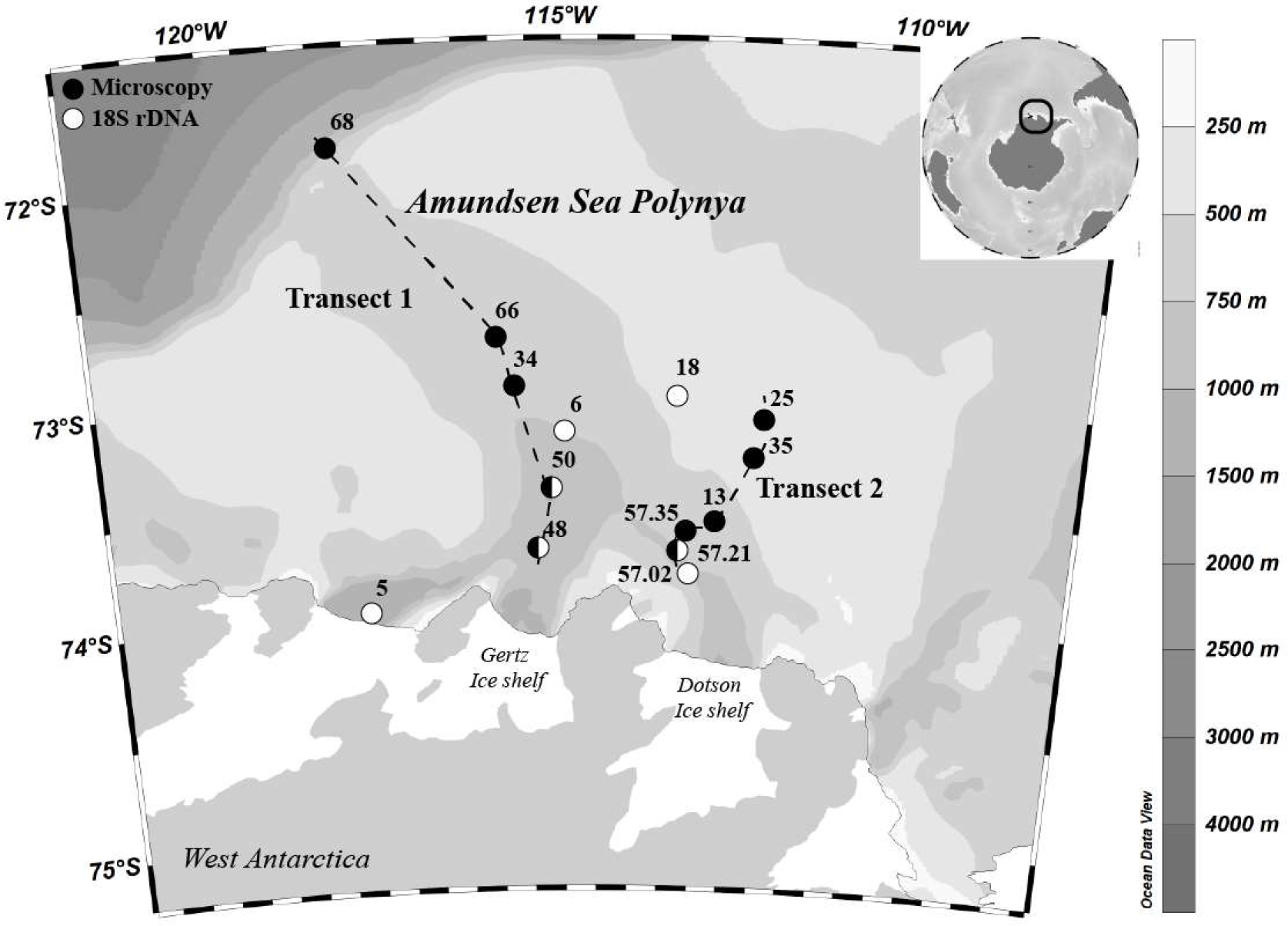
Location of the ASPIRE stations, sampled for microscopy analyses (•) and 18S rDNA (○). Depth contours are illustrated in gray scale. Station numbers are indicated.

### 2.2. Microzooplankton community and biomass

Microscopy samples were collected at 10 of the 14 microzooplankton sampling stations (Fig. 1) traversing the ASP from the surface (2 - 5 m), depth of fluorescence max. (DFM, 10 - 40 m), and below the fluorescence peak (50 - 180 m). Water from the Niskin bottles was gently siphoned through silicon tubes into 300 ml amber colored glass bottles and fixed in acidic Lugol’s solution (2% final concentration). Bottles were stored cool and dark (max. 12 months) until appropriate sized sub-samples (depending on cell concentration) were transferred into sedimentation chambers, allowed to settle for 24 hours and microplankton identified and counted under microscope at the Latvian Institute of Aquatic Ecology. Cell volumes were calculated using appropriate geometric shapes following (Olenina, 2006). To correct for shrinkage due to Lugol preservation, cell volumes were adjusted by a factor of 1.33 (Stoecker et al., 1994). Biomasses of dinoflagellates, loricate and aloricate ciliates were calculated using carbon conversion factors by Menden-Deuer and Lessard (2000).

### 2.3. RNA extraction, sequencing and taxonomic identification

Between 1-6 liters of seawater were pre-filtered through a 20μm sieve and then sequentially filtered through 3 polycarbonate filters. Filters were flash-frozen and stored at −80° C. RNA was extracted using the NucleoSpin® RNA L kit (Macherey-Nagel) and quantified using a Nanodrop ND-1000 Spectrophotometer. To remove DNA from RNA extracts, we used the TurboDNA kit (Ambion). RNA were reverse transcribed using the RT Superscript III random primers kit (Invitrogen) and hypervariable V4 region was amplified using the universal primers TAReuk454FWD1 (5’-CCAGCASCYGCGGTAATTCC-3’) and TAReukREV3 (5’-ACTTTCGTTCTTGATYRA-3’) (Stoeck et al., 2010). Triplicate amplicon reactions were pooled and purified using NucleoSpin® Extract II (Macherey-Nagel). Purified amplicons were quantified with Picogreen Invitrogen) and pooled in equimolar concentration. Amplicon sequencing was carried out on a *454* GS FLX Titanium system (*454* Life Sciences, USA) at Genoscope (http://www.genoscope.cns.fr/spip/, France).

All reads were were processed with Quantitative Insight Into Microbial Ecology pipeline, QIIME v1.4 (Caporaso et al., 2010). Only reads between 200-500 bp were used. Reads were quality controlled and denoised using DeNoiser v 0.851 (Reeder & Knight, 2010) implemented in QIIME. Subsequently, reads were clustered into Operational Taxonomic Units (OTUs) using UCLUST v1.2.22 (Edgar, 2010) at 99% similarity. Chimeras were detected and removed using ChimeraSlayer with a reference database derived from PR2 (Guillou et al., 2013). Representative reads were assigned to taxonomy by BLASTing them against the databases SILVA v108 (Quast et al., 2013), the PR2. Sequences are publicly available at ENA (PRJEB23910). Only OTUs representing > 0.1 % of the total relative abundance of ciliates and dinoflagellates were further studied. Maximum likelihood trees were computed with MEGA7 (Kumar et al., 2016).

### 2.4. Induced bloom microcosm experiment set-up

The succession of the ciliate and dinoflagellate community was followed during the course of a 15 day *Phaeocystis* bloom induced in St. 35 DFM water (12 m). Triplicates incubations in 12 l collapsible plastic containers were carried out for both unfiltered water and 200 μm filtrates. The water from the Niskin bottles was gently siphoned into a 60 l bucket, and vitamins and nutrients (15 μM NH_4_Cl and 1 μM Na_2_HPO4) were added before carefully mixing the water for the triplicate aliquots. The filtered treatment was done to ensure absence of metazoan predators of dinoflagellates and ciliates, by gentle reverse filtration through 200 μm mesh size filters. The containers were incubated at *in-situ* light (PAR = 2.04 μmol m^−2^ s^−1^) and temperature (0 ± 0.5 °C) for 15 days and mixed by gentle rotation of the containers every 8 h. The light and temperature conditions were monitored regularly during the experiment. Samples for nutrients, chlorophyll *a* (chl *a*), microzooplankton biomass and species composition (analyzed following same procedure as for *in situ* samples) were collected at the start of the incubation and after 2, 4, 7, 10, 13, and 15 days. HNF were also measured by flow cytometry according to Christaki et al. (2011), while bacteria were enumerated with the flow cytometry method described by Gasol and del Giorgio (2000) for bacteria. Samples for 18S rRNA based community analysis as well as, *Phaeocystis* and diatom abundances were collected at the start, day 7 and at the end of the experiment (pooled triplicates). In order to avoid disturbance from air bubbles during mixing, air was squeezed out of the collapsible container after every sampling. All containers and materials were acid washed prior to use.

Fifty ml samples for phosphate, nitrate, ammonium and silicate were measured onboard, on a 5 channels auto-analyzer (Lachat Instrument QuikChem FIA +8000 serie). For chl *a* measurements, triplicate 50 ml aliquots from each container were filtered onto Whatmann GF/F filters (0.7 μm) and extracted in 96% ethanol for 12 - 24 h (Jespersen & Christoffersen, 1987) before being analyzed on a fluorometer (Turner Model 10AU), both before and after acidification (Yentsch & Menzel, 1963). Growth rates were calculated for each time step using the equation:

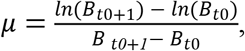
where B is the concentration or biomass at sampling day t_0_ and at the following sampling day t_0+1_.

### 2.5. Data analysis

Station map and physical conditions and distributions of chl *a* and microzooplankton along two transects was visualized using the Ocean Data View (weighted-average gridding, v. 4.7.1). Microzooplankton species richness (Margalef D), diversity (Shannon-Wiener H’) and evenness (Pielou’s J’) were calculated using the equations:

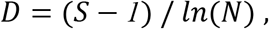

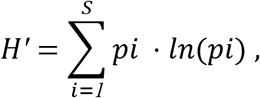

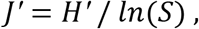
where S is the no. of species, N is the no. of individuals and p_i_ is the proportion of the *i*th species. Size evenness was calculated using cell abundances within five biovolume size groups (<2000, 2000-10000, 10000-50000, 50000-250000, >250000 μm^3^).

Spearman rank correlation analysis between microzooplankton community indices and environmental parameters was carried out in Sigmaplot v. 12 (Systat Software, Inc.). Correlation between Bray-Curtis resemblance matrices of microzooplankton species biomass and Euclidian distance resemblance matrices of environmental parameters was tested using the BEST analysis in Primer v. 6.1.7 (Primer-E, Ltd). The BIOENV algorithm and Spearman rank correlation method was used. A stepwise distance-based linear model permutation test (DistLM, McArdle & Anderson, 2001) was also performed to identify environmental variables best predicting community variation. The stepwise routine was run employing 9,999 permutations and using the AICc (Akaike’s information criterion with second order correction) selection criterion. Results were visualized with a distance-based redundancy analysis (dbRDA, Anderson et al., 2008). Analysis of similarities in community composition (ANOSIM) was carried out by pairwise testing between different depth strata. To identify the species contributing most to the similarity between samples, a SIMPER analysis (SIMilarity PERcentage, Clarke & Warwick, 2001) was performed. Lastly, testing of similarities between microscopy and 18S rRNA Bray-Curtis resemblance matrices was done by RELATE analysis of Spearman rank correlations. All biomass and environmental data were log transformed while abundance data was fourth root transformed. Environmental data was normalized by subtracting means and dividing by the standard deviation (z-scores).

## 3. Results

### 3.1. Hydrography and microzooplankton

The upper part of the water column (30 - 50 m) was characterized by increasing temperatures with increasing latitude and proximity to the ice shelf (Fig. 2). Inversely, salinity increased with decreasing latitude and proximity to the sea ice surrounding the polynya. Two stations along transect 2 deviated from this pattern; St. 57.21 where the upper ~100 m was mixed as it was located in the wake of a drifting iceberg (Randall-Goodwin et al., 2015; Dinasquet et al., 2017), and St. 35 characterized by high surface temperatures. Detailed information on the hydrography is presented elsewhere (Randall-Goodwin et al., 2015; Yager et al., 2016).

**Figure 2:**
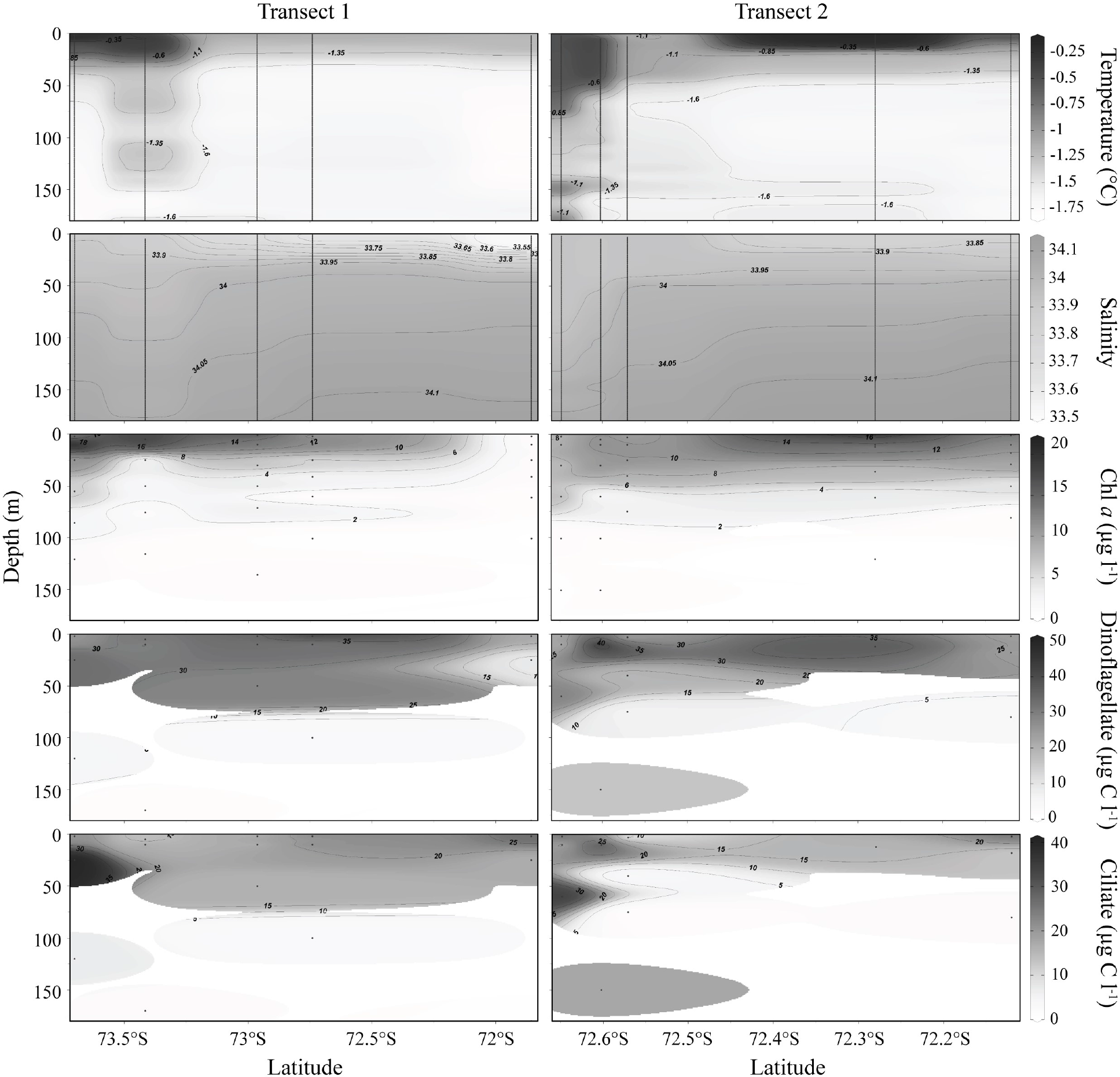
Vertical distribution patterns of temperature, salinity, chl *a*, dinoflagellate and ciliate biomass along two latitudinal transects crossing the polynya. Data also presented in Yager et al. (2016).

Chlorophyll *a* (Chl *a*) concentration was highest in the top 40 m of the water column and where temperatures were elevated, with St. 57.21 and 57.35 as exceptions (Fig. 2). *Phaeocystis antarctica* likely represented most of the phytoplankton contribution to Chl *a* followed by diatom (Table S1, Alderkamp et al., 2015; Yager et al., 2016). Dinoflagellates and ciliates made up most of the microzooplankton biomass (Table S1). Subsurface peaks in dinoflagellate and particularly ciliate biomass occurred below the chl *a* max. at stations closer to the ice shelf and in the wake of the drifting iceberg. Dinoflagellate biomass was also high at St. 35, while elevated surface ciliate biomass occurred at St. 68 at the fringe of the polynya.

### 3.2. Microzooplankton community patterns and environment

The correlation between environmental parameters and microzooplankton community was tested both against indices of community structure (species richness, diversity, eveness) and composition (Bray-Curtis dissimilarity). Overall, ciliate biomass was correlated to most environmental variables measured, while dinoflagellate biomass and evenness in microzooplankton size distribution was correlated to all (Table 1). Depth was likely a main factor driving community structure (based on microscopy), as negative correlations were observed with variables that generally increase with depth (salinity, dissolved inorganic nitrogen DIN), and positive correlations with variables decreasing with depth. Species richness and diversity displayed similar patterns and were generally positively correlated to heterotrophic nanoflagellate (HNF) abundance. Species richness also increased with chl *a* and concentration of pigment markers for diatoms (fucoxanthin) and prymnesiophytes, such as *Phaeocystis* spp. (19’-hexanoyloxyfucoxanthin).

**Table 1:** Spearman correlation coefficients between environmental variables and dinoflagellate and ciliate species richness (D), diversity (H'), evenness (J' species), size distribution evenness (J' size) and biomasses.

|  | **D** | **H'** | **J' species** | **J' size** | **Dinoflagellates** | **Ciliates** |
| --- | --- | --- | --- | --- | --- | --- |
| **Depth** | 0.105 | -0.179 | 0.082 | -0.535** | -0.585** | -0.431* |
| **Temperature** | 0.284 | 0.155 | -0.114 | 0.553** | 0.682** | 0.609** |
| **Salinity** | -0.422* | -0.47* | -0.143 | -0.640** | -0.714** | -0.544** |
| **PAR<sup>1</sup>** | 0.105 | 0.212 | 0.038 | 0.469* | 0.515** | 0.461** |
| **DIN<sup>2</sup>** | -0.497** | -0.331 | 0.005 | -0.692** | -0.825** | -0.460* |
| **Chl *a*** | 0.402* | 0.291 | -0.035 | 0.738** | 0.782** | 0.337* |
| **Fucoxanthin** | 0.464* | 0.350 | 0.026 | 0.820** | 0.829** | 0.477* |
| **19'-Hexanoyloxyfucoxanthin** | 0.474* | 0.283 | -0.049 | 0.723** | 0.756** | 0.268 |
| **HNF<sup>3</sup>** | 0.540** | 0.502** | 0.155 | 0.773** | 0.842** | 0.576** |
| **Bacteria** | 0.206 | 0.117 | -0.169 | 0.416* | 0.556** | 0.012 |

\* $p < 0.05$ , \*\* $p < 0.01$ , <sup>1</sup>Phyotosynthetically Active Radiation, <sup>2</sup>Dissolved Inorganic Nitrogen, <sup>3</sup>Heterotrophic Nanoflagellates

The environmental variables that best explained the microzooplankton community composition patterns were depth, bacterial and HNF abundances, and fucoxanthin concentration (BEST, Rho = 0.72, p = 0.01). Distance based linear models (DistLM) found microzooplankton community composition to be related to all environmental parameters tested (depth, temperature, salinity, density, PAR, total DIN, chl *a*, bacterial and HNF abundance, 19’-hexanoyloxyfucoxanthin and fucoxanthin concentration, p < 0.005). HNF and bacterial abundance generated the lowest AICc score thus explaining most of the variance in microzooplankton community composition (AICc = 192.81, R^2^ = 0.38, Fig. 3).

**Figure 3:**
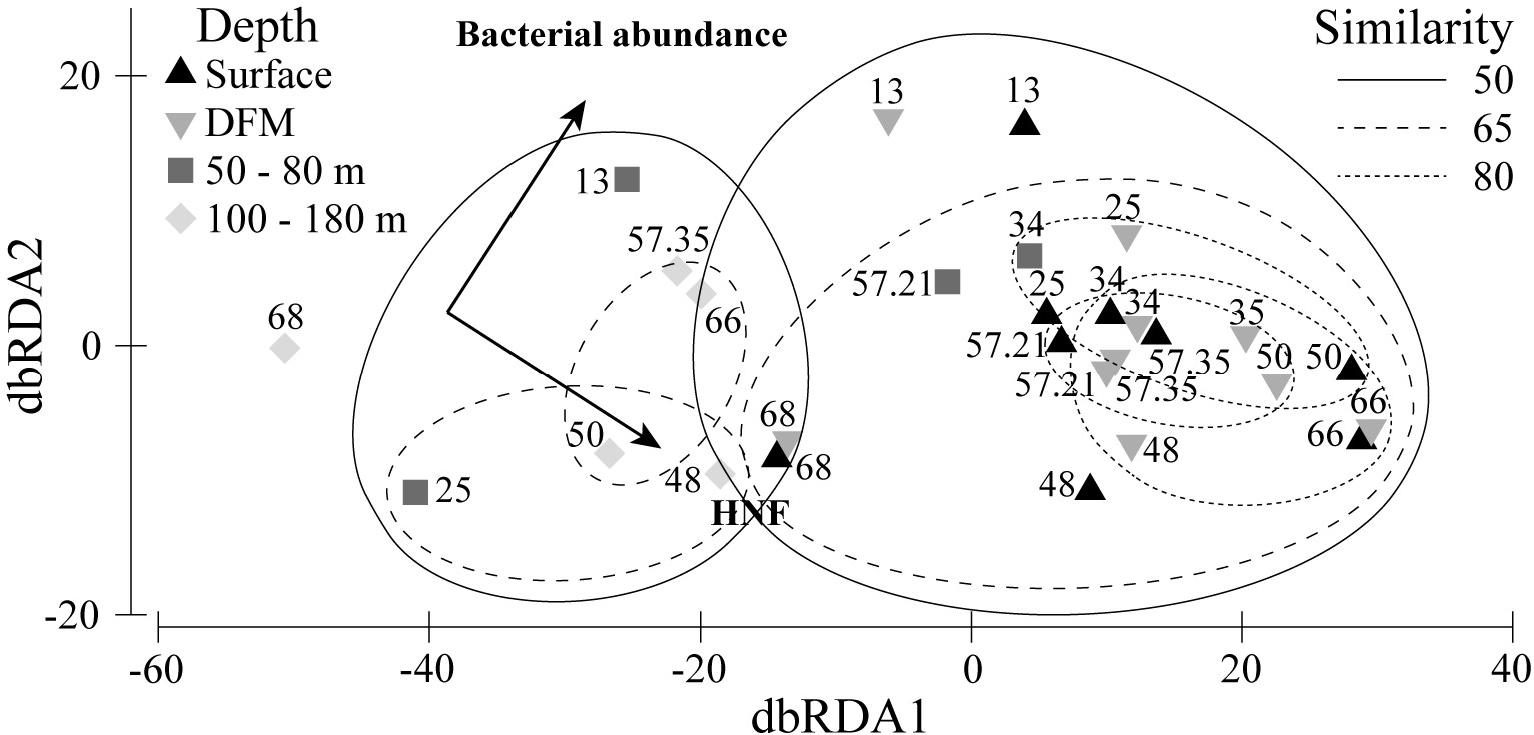
Distance based linear modelling (DistLM) plot on Bray-Curtis similarities of log transformed ciliate and dinoflagellate species biomass data at 28 sampling points in relation to log transformed and normalized environmental parameters. dbRDA1 explained 88.3% of fitted and 34% of total variation while dbRDA2 explained 11.7% and 4.5%, respectively. Overlaid is community similarity levels based on Bray-Curtis resemblance matrices. Station numbers are indicated.

### 3.3. Taxonomy and distribution

Microscopy-based microzooplankton community dissimilarities were greater between the surface and depth of fluorescence max. (DFM) than between sampling stations (p = 0.12), but differed significantly from the deeper samples (p < 0.03, Fig. 3). Station 68 located at the edge of the polynya and St. 13, which had the lowest species richness and diversity of all stations (data not shown), displayed the least similarity to the communities seen at the other stations (Fig. 3).

Microscopy counts indicated that the community was dominated by dinoflagellates constituting on average 58% of the microzooplankton biomass (Fig. 4, Table S1). *Gymnodinium* spp. and heterotrophic *Gyrodinium spirale* and *Protoperidinium* spp. (mainly *P. depressum*) accounted for 87% of the dinoflagellate biomass, but differed in distribution patterns (Fig. 4A). *Gyrodinium* spp. tended to decrease with distance to the ice shelf, while *G. spirale* and *Protoperidinium* spp. showed inverse distributions between transects 1 and 2. The ciliate community was dominated by heterotrophic tintinnids, *Strombilidium* spp. (mainly *S. epacrum* and *S. spiralis*) and mixotrophic *Strombidium* spp. that together contributed with 91% of the ciliates biomass (Fig. 4B). Along transect 1 tintinnid and *Strombidium* spp. biomass increased with distance to the ice shelf while *Strombilidium* spp. tended to decrease. However, on transect 2 both tintinnids and *Strombilidium* spp. represented the highest biomass on St. 57.35. The heterotrophic *Didinium nasutum* represented the highest biomass only on station 48. Station 13 and 57.21 were characteristic in featuring fewer microzooplankton than adjacent stations, and the ciliate community at St. 13 was dominated solely by tintinnids (Fig. 4).

**Figure 4:**
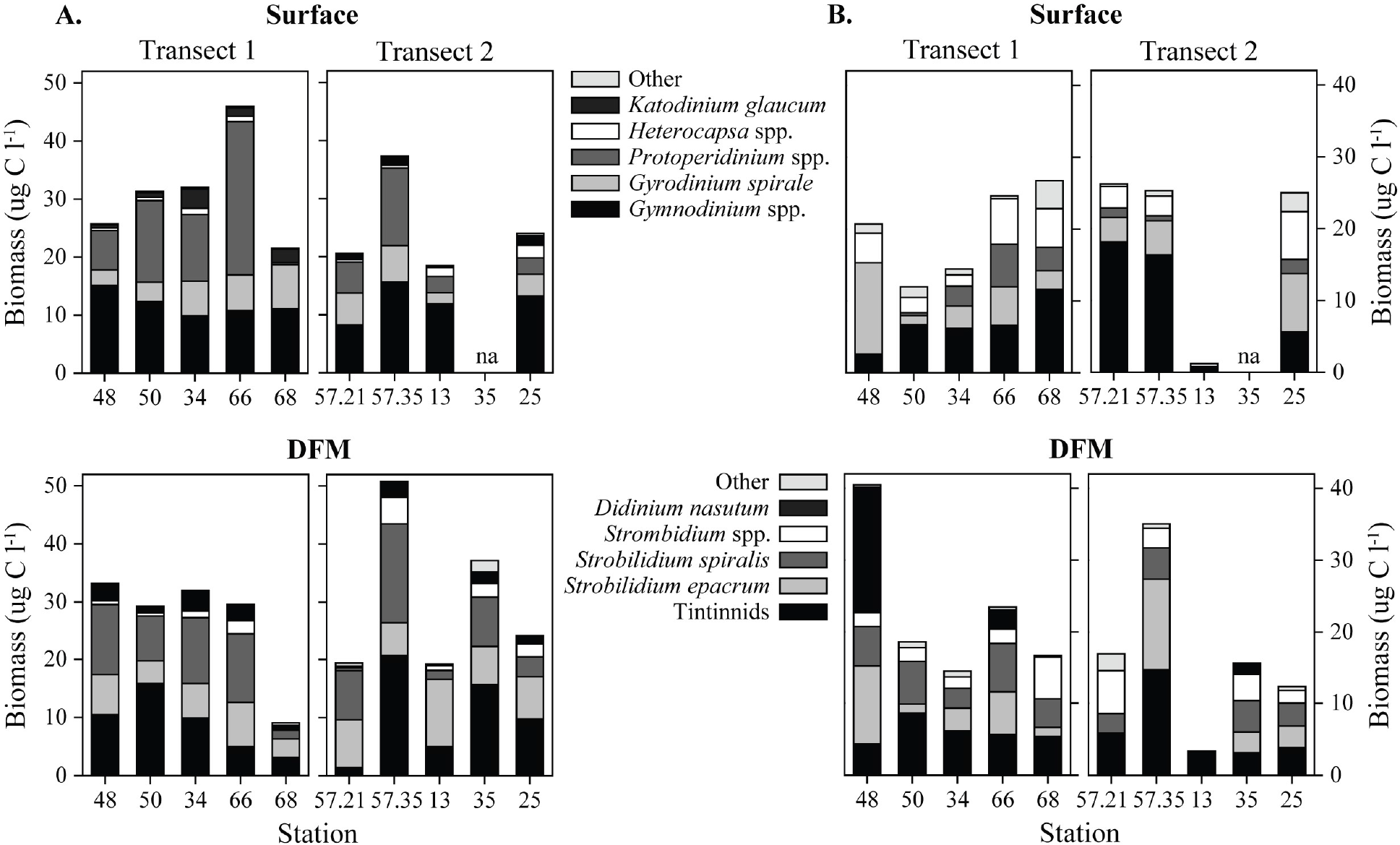
Biomass distribution of dominant dinoflagellate (A) and ciliate (B) taxa along two transects at the surface (2 - 5 m) and depth of fluorescence max. (DFM, 10 - 40 m).

The 18S rRNA analysis confirmed that the microzooplankton community was dominated by dinoflagellates representing on average 85% of the microzooplankton amplicons at surface and DFM and 68% in deeper waters (>400 m, Fig. S1). The main OTU found in all samples was related to the dinoflagellate SL163A10 (99% identity) closely related to Gymnodiniales (Fig. 5A). Other important dinoflagellates were closely related to Peridiniales increasing with depth and other Gymnodiniales related to *Gymnodinium* sp. and *Gyrodinium* sp. (Fig. 5A, S1). Ciliates represented on average 15% of the microzooplankton related OTUs at surface and DFM and their contribution increased to ~ 32% atdepth. The dominant ciliate OTUs were closely related to potential parasites from the Ventrata order (Fig. 5B, S1). Other abundant ciliate OTUs were related to Choreotrichia, Oligotrichia, and Haptoria.

**Figure 5:**
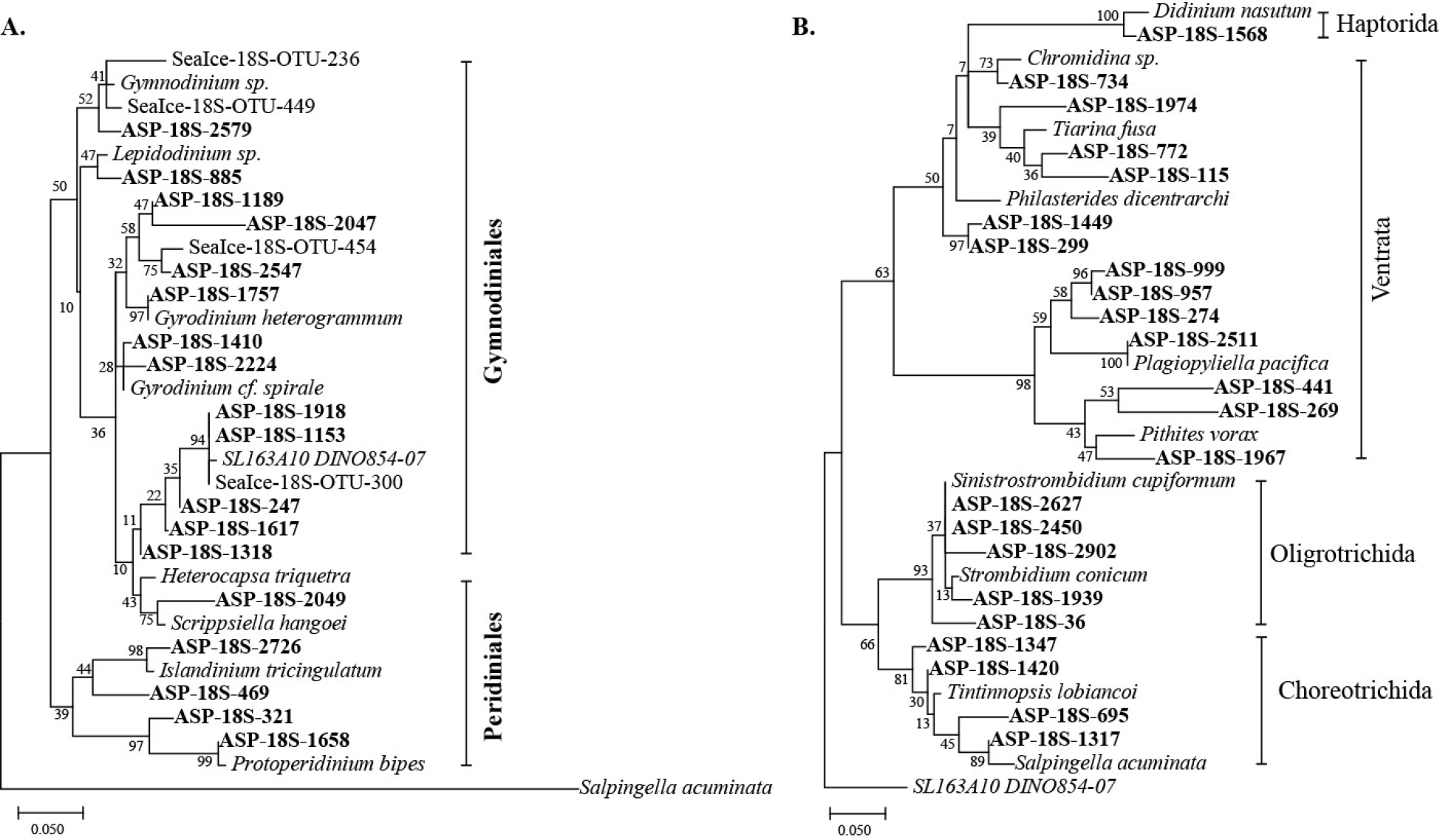
Maximum likelihood tree of the operational taxonomic units (OTU) closely related to dinoflagellates (A) and ciliates (B). Only OTUs representing more than 0.1% of the total relative abundance of ciliates and dinoflagellates 18S RNA gene reads are included. Reference sequences are indicated in italics. Sea ice OTUs come from a study of protists in the ASP sea ice during the same sampling time (Torstensson et al. 2015). Bootstrap values (n = 1000) are indicated at nodes; scale bar represents changes per positions.

Resemblance analysis of 12 microzooplankton taxonomic orders identified in either the sequencing or microscopy dataset (Fig. 4, S1, Table S1) from 6 overlapping samplings (St. 48, 50, 57.21 at surface and DFM, Fig. 1) showed no significant correlation between their relative abundance in the 18S rRNA amplicon dataset and biomass (Rho = 0.46, p = 0.11) or cell abundance (Rho = 0.4, p = 0.08).

### 3.4. Succession during an induced Phaeocystis bloom

During the course of the induced experimental *Phaeocystis* bloom, no major differences were observed between filtered and unfiltered water. No metazoans were observed during the experiment in any treatment. Chl *a* increased about 5-folds in both treatments over the course of the incubation (Fig. 6A). The increase in chl *a* was mainly associated with a 4-fold increase in *Phaeocystis antarctica* reached an average biomass of 750 μg C l^−1^ on day 15. Many large *P. antarctica* colonies were observed in all replicate containers. Diatoms also increased up to an estimated biomass of 250 μg C l^−1^ with a faster growth rate than *P. antarctica* (Table S2). The dominant dinoflagellates *Gymnodinium* spp. and *Gyrodinium spirale* increased over the course of the bloom, especially *Gymnodinium* spp., with a growth rate of ~0.18 d^−1^, reaching a biomass of 100 μg C l^−1^ in the unfiltered treatments (Fig. 6.B, C, D, Table S2). The most abundant dinoflagellate OTU was related to SL163A10 (data not shown). Other dinoflagellate taxa remained stable or decreased during the experiment (Fig. 6D, Table S2). Tintinnids were the main group of ciliates responding to the extended bloom with a biomass of up to 8 μg C l^−1^ and a growth rate of 0.21 d^−1^ in the unfiltered treatment (Fig. 6E, F, Table S2). Other ciliates (e.g. *Strobilidium* spp. and *Strombidium* spp.) decreased slightly during the course of the experiment.

**Figure 6:**
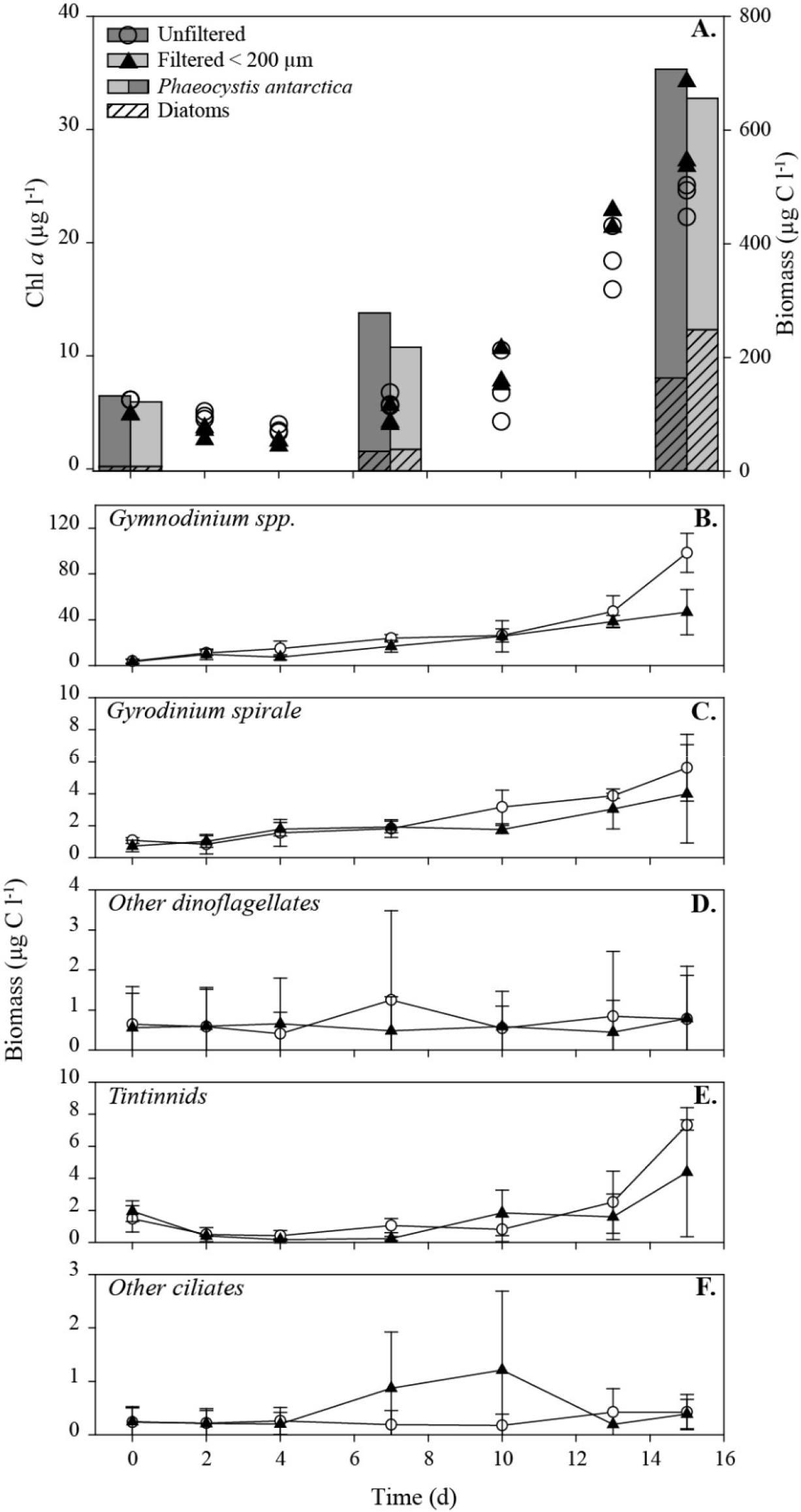
Chlorophyll *a* concentration and biomass of dominant phytoplankton (A). Biomass of the dominant dinoflagellates (*Gymnodinium* spp. B., *Gyrodinium spirale* C. and others D.) and ciliates (Tintinnids E. and others F.) over the course of the induced *Phaeocystis antarctica* bloom. Mean ± SD for 3 replicates for each treatment.

## 4. Discussion

In early summer, the open waters of the Amundsen Sea polynya (ASP) harbor extensive episodic blooms of the colony forming prymnesiophyte *Phaeocystis* sp. (Alderkamp et al., 2012; Arrigo et al., 2012). At the time of sampling in summer 2010-2011, the full net primary production was principally exported to deeper water or grazed by microzooplankton (Yager et al., 2016). Our study suggests that grazing pressure on the bloom in the ASP was mainly due to dinoflagellates known to be mixotrophs and heterotrophs, and hence our work provides further insights about the composition and dynamics of the microzooplankton community in the poorly explored polynyas of the Southern Ocean (SO).

### 4.1. Patterns in microzooplankton community structure

Known heterotrophic and mixotrophic dinoflagellates, mainly *Gymnodinium*, *Gyrodinium* and *Protoperidinium,* dominated the ASP microzooplankton community. These three genera were also dominant in the ASP in time periods following after the present study (Yang et al., 2016) and are known to be widely distributed across the SO (Stoecker et al., 1995; Safi et al., 2007; Pearce et al., 2008; Garzio & Steinberg, 2013; Christaki et al., 2015). Less abundant ciliates known to be heterotrophic or mixotrophic, particularly loricate tintinnids and aloricate *Strobilidium* spp. and *Strombidium* spp., also constituted an important part of the community. This differs from other studies in the Amundsen Sea (AS) where Oligotrichs; mainly *Strombidium* spp. and *Tontonia* spp., and the Choreotrich; *Lohmanniella oviformis* dominated the ciliate assemblage while tintinnids were few and *Strobilidium* spp. was absent (Dolan et al., 2013; Jiang et al., 2014; Jiang et al., 2016). *Strobilidium* and *Lohmanniella* are morphologically similar and may have been misidentified in either our study or by (Jiang et al., 2014; Jiang et al., 2016). High abundance of both genera have been reported from the AS (Wickham et al., 2011) and the Kerguelen area in the SO (Christaki et al., 2015).

Microzooplankton community structure changed with depth and distance to the ice shelf, and positively correlated mainly with abundance of heterotrophic nanoflagellates (HNF), bacteria and concentration of the diatom pigment marker fucoxanthin. These relationships may be explained by all trophic levels responding to the same environmental drivers or by direct predator-prey relationships. Low picophytoplankton biomass in the ASP (Lee et al., 2012; Yang et al., 2016) may have resulted in a dietary shift for HNF to primarily graze on bacteria (Gonzalez et al., 1990; Pearce et al., 2011) and in turn being grazed by heterotrophic dinoflagellates and ciliates (Kuparinen & Bjornsen, 1992; Jürgens et al., 1996). Some ciliates are also capable of grazing directly on bacteria (Sherr & Sherr, 2002). Larger diatoms (>15 μm in length), which accounted for most of the diatom biomass (data not shown), are mainly grazed by heterotrophic dinoflagellates (Hansen, 1992; Sherr & Sherr, 2007; Grattepanche et al., 2011). Thus, changes in diatom contribution to a *Phaeocystis* dominated phytoplankton community, as well as HNF abundance are expected to propagate into the microzooplankton community structure. Although mesozooplankton biomass was generally low, predation likely affected the microzooplankton assemblage at St. 13, which experienced the highest mesozooplankton grazing (Wilson et al., 2015; Yager et al., 2016)and where the ciliates were almost exclusively tintinnids, a group known to be more resistant to metazoan predation (Stoecker, 2012). Lastly, upwelling of deeper water masses downstream of a drifting iceberg (St. 57.21) and contribution of sea ice melt water (St. 68) (Randall-Goodwin et al., 2015; Dinasquet et al., 2017) was seen to affect biomass and community structure.

### 4.2. Morphological and molecular analyses of microzooplankton composition and dynamics

While morphological and molecular information on microzooplankton are generally not directly comparable (Medinger et al., 2010; Monchy et al., 2012; Christaki et al., 2015) due to large variations in 18S rDNA copies per organism (Zhu et al., 2005; Gong et al., 2013) and the limited taxonomic resolution provided by morphology, the two methods are complementary.

Both methods showed that the microzooplankton community was dominated by dinoflagellates, in particular the Gymnodiniaceae family dominated at most stations. Surprisingly, Peridiniales-related sequences were few despite representing a substantial fraction of the dinoflagellate biomass, possibly due to relatively inefficient RNA extraction from thecate dinoflagellates. Members of the Gymnodicianiceae family have very similar morphological attributes, which makes them difficult to distinguish (Gast et al., 2006). Here, through sequencing, the dominant dinoflagellates were identified as closely related to SL163A10; a species also very abundant in the Ross Sea Polynya (Gast et al., 2006) and the Antarctic peninsula (Luria et al., 2014). This dinoflagellate was also found to be the dominant protist in the ASP sea ice at the time of sampling, where it may play an important ecological role (Torstensson et al., 2015).

Whereas both methods reported similar proportions of ciliates and dinoflagellates, the detailed taxonomic information was not comparable. For instance, *Strombidium* spp., *Strobilidium* spp. and tintinnids had the highest biomasses, but the sequencing did not match their relative biomass or abundance contribution. Tintinnids, which are detectable by both methods (Bachy et al., 2011), were underrepresented in our sequencing dataset. Tintinnids, *Strombidium* spp. and *Strobilidium* spp. have distinct morphological features and it is unlikely that they were misidentified through microscopy. On the other hand, ciliates related to the Oligohymenophorea class were the most abundant in the sequencing dataset especially in deep waters, as found in other studies (Zoccarato et al., 2016; Zhao et al., 2017). They may have been overlooked in the microscope as their pelagic stage is usually a dormant cyst like form, which is difficult to identify. Interestingly, many taxa related to Oligohymenophorea are potential symbionts and parasites of crustaceans (Gómez-Gutiérrez et al., 2006; Gomez-Gutierrez et al., 2012), but their importance for zooplankton population dynamics in the Southern Ocean is so far unknown.

### 4.3. Microzooplankton ecology during a Phaeocystis bloom in the ASP

*Phaeocystis antarctica* was by far the most abundant phytoplankton in the ASP at the time of sampling (Alderkamp et al., 2015; Yager et al., 2016, Table S1). Particular interest have been given to *Phaeocystis* due to its production of DMSP (reviewed in Liss et al., 1994) and high rates of primary production (DiTullio et al., 2000; Alderkamp et al., 2012). Different species of this prymnesiophyte are ubiquitous in marine environments where they can form dense blooms (Schoemann et al., 2005) has a global distribution and is found as different species in very different marine environments. Rapid increase of *Phaeocystis* may in part be ascribed to its capacity to form large colonies not readily grazed on by micro- and mesozooplankton (Caron et al., 2000; Jakobsen & Tang, 2002; Nejstgaard et al., 2007; Grattepanche et al., 2011). The capacity of some microzooplankton species, such as *Gyrodinium* spp., *Gymnodinium* spp. and tintinids, to graze on single cells and small colonies of *Phaeocystis* (Admiraal & Venekamp, 1986; Bjørnsen & Kuparinen, 1991; Stoecker et al., 1995; Nejstgaard et al., 2007; Grattepanche et al., 2010), would nevertheless explain the dominance of these species at the time of sampling. *Gyrodinium* spp., *Gymnodinium* spp. and tintinids were also found to co-dominate during *Phaeocystis* blooms in McMurdo Sound and the North Sea (Weisse & Scheffel-Möser, 1990; Stoecker et al., 1995). The intense phytoplankton growth observed in the induced bloom experiment, despite an increased dominance of known *Phaeocystis* grazers, suggested that *P. antarctica* were not controlled by microzooplankton herbivory possibly because of low grazing rates (Caron et al., 2000; Yager et al., 2016) at this colonial stage of the bloom. However, higher grazing rates of 90% d^−1^ were observed later in the ASP (Yang et al., 2016), suggesting the capacity of microzooplankton to control the later stage of the *Phaeocystis* bloom where colonies breakup into single cells. Single *Phaeocystis* cells are more vulnerable to predation and are often released from colonies later in the season possibly when nutrients become limiting (Jakobsen & Tang, 2002; Smith et al., 2003; Nejstgaard et al., 2007).

Most of the ciliates we observed in the ASP were heterotrophs with the ability to graze on single cell *Phaeocystis* and small diatoms (Grattepanche et al., 2010; Dolan et al., 2013). The dominant dinoflagellate *Gymnodinium* spp. is capable of grazing on small *Phaeocystis* colonies (Grattepanche et al., 2011). In the present study, the most abundant microzooplankton Gymnodiniaceae SL163A10 engage in kleptoplasty of *P. antarctica* chloroplasts (Gast et al., 2007). This dinoflagellate is ubiquitous in the SO, in waters and sea ice (Gast et al., 2006; Luria et al., 2014; Torstensson et al., 2015, this study), suggesting that its mixotrophic life strategy is highly successful. The observed microzooplankton community shift towards the dominance of *Gymnodinium* spp. (83% of microzooplankton biomass) in the microcosm experiment also underlines this species ability to thrive in an environment dominated by *Phaeoystis* colonies. Nevertheless, the relative importance of its primary vs. secondary production within the food web is not known. Mixotrophic dinoflagellates are important phytoplankton grazers in the SO open waters and semi-enclosed polynyas (Gast et al., 2006; Christaki et al., 2015) and potentially important prey for zooplankton grazers, although they did not sustain high biomass of zooplankton in the ASP (Lee et al., 2013; Wilson et al., 2015).

### 4.4 Concluding remarks

During this study, colonial *Phaeocystis* were grazed at low rates, but sustained a high biomass of specialized microzooplankton capable of grazing on them. Although taxa known to be mixotrophic were important, heterotrophy appeared to be the main life strategy for the microzooplankton. The presence of *Phaeocystis* colonies appeared to determine the key microzooplankton taxa, while other potential prey seemed more important for shaping the community composition of less abundant taxa within the ASP. The early shift in community composition observed during the induced bloom experiment as well as the major differences in microzooplankton community composition and biomass observed few days later in the polynya (Jiang et al., 2014; Yang et al., 2016) are consistent with the pronounced and selective impact of *Phaeocystis* blooms on growth, biomass and composition of the cooccurring microzooplankton. Whereas these interactions undoubtedly affect biogeochemical nutrient fluxes mediated by the microbial loop in euphotic Southern Ocean waters, the consequences for vertical carbon export remains to be addressed.

## Acknowledgments

We gratefully acknowledge the support from captain and crew of the *RVIB Nathaniel B. Palmer*, the Raytheon support team onboard and our chief scientists P. Yager. We also thank K. Arrigo’s team for sharing pigment data. *Oden Southern Ocean*, SWEDARP 2010/11, was organized by the Swedish Polar Research Secretariat and National Science Foundation Office of Polar Programs. This work was supported by the Swedish Research Council [grant 2008-6430] to S. Bertilsson and L. Riemann and [grant 824-2008-6429] to P.-O. Moksnes and J. Havenhand) and by the US National Science Foundation through the ASPIRE project [NSF OPP-0839069] to P. Yager.

## Declarations of interest

none

